# Selective JAK Inhibition Reveals Paradoxical and Hierarchical Control of interferon-γ-driven Autoimmunity in AIRE Deficiency

**DOI:** 10.64898/2026.03.05.709894

**Authors:** Eliezer Heller, Lucas dos Santos Dias, Michail S. Lionakis

**Affiliations:** Fungal Pathogenesis Section, Laboratory of Clinical Immunology and Microbiology, National Institute of Allergy and Infectious Diseases, National Institutes of Health, Bethesda, Maryland, USA

## Abstract

Autoimmune polyendocrinopathy-candidiasis-ectodermal dystrophy (APECED) is caused by impaired central immune tolerance due to deficiency of the Autoimmune Regulator (AIRE) and is characterized by severe, multiorgan autoimmunity. We recently identified interferon-γ (IFN-γ) as a dominant driver of immunopathology in APECED and showed that treatment with the JAK1/2 inhibitor ruxolitinib ameliorates disease in both AIRE-deficient mice and patients. However, broad JAK inhibition is associated with clinically relevant toxicities, raising the question of whether selective targeting of individual JAK pathways can retain efficacy while sparing nonpathogenic immune programs. Here, we systematically evaluated the effects of selective JAK1, JAK2, and JAK3 inhibition in *Air*e^−/−^ mice. Selective JAK1 and JAK2 inhibition reduced autoimmune tissue injury, suppressed IFN-γ signaling, and decreased accumulation of pathogenic T cells, with JAK2 inhibition providing the most robust protection, comparable to ruxolitinib. In contrast, selective JAK3 inhibition decreased T cell accumulation, but paradoxically increased the proportion of IFN-γ-producing T cells and did not significantly attenuate IFN-γ-driven tissue inflammation. These findings reveal an unexpected uncoupling between lymphocyte burden and pathogenic cytokine bias and identify IFN-γ signaling as hierarchically dominant over γc-dependent pathways in AIRE deficiency. Together, our data indicate that effective control of APECED-associated autoimmunity requires direct suppression of the IFN-γ-JAK2 axis rather than generalized lymphocyte inhibition and suggest that selective JAK2 targeting may represent a rational strategy to preserve therapeutic efficacy while minimizing disruption of JAK1-and γc-dependent immune functions.

## Introduction

Autoimmune polyendocrinopathy-candidiasis-ectodermal dystrophy (APECED), also known as Autoimmune polyendocrine syndrome type-1 (APS-1), is a life-threatening multiorgan autoimmune disease caused by inherited deficiency of the autoimmune regulator (AIRE) that impairs central immune tolerance [1–11]. We recently demonstrated that excessive interferon-γ (IFN-γ) signaling is a dominant driver of immunopathology in APECED and that inhibition of IFN-γ responses with the FDA-approved JAK1/2 inhibitor ruxolitinib ameliorates tissue injury in both *Aire*^−/−^ mice and APECED patients [12, 13]. These findings established IFN-γ-JAK-STAT signaling as a therapeutically actionable pathway in AIRE deficiency but also raised questions about the necessity and consequences of broad JAK inhibition.

In parallel, the emergence and regulatory approval of selective JAK1, JAK2, and JAK3 inhibitors has created new opportunities to interrogate and therapeutically target JAK pathways while sparing nonpathogenic immune programs. Preclinical studies in IFN-γ-driven inflammatory settings, including alopecia areata, have shown that selective inhibition of individual JAKs can ameliorate disease; notably, in the skin, JAK3 inhibition has been most effective, followed by JAK1 inhibition, with comparatively limited effects of JAK2 inhibition on IFN-γ production and hair regrowth [14]. At the same time, long-term nonselective JAK inhibition has been associated with clinically important toxicities, including increased susceptibility to viral infections, disruption of lymphocyte homeostasis, interference with growth hormone, leptin, erythropoietin, and thrombopoietin receptor signaling, thrombosis, and squamous cell carcinoma [15, 16]. These considerations make it critical to define whether selective inhibition of specific JAKs can phenocopy the protective effects of ruxolitinib while minimizing collateral immune disruption.

Here, we systematically evaluated selective JAK1, JAK2, and JAK3 inhibitors and compared their effects with ruxolitinib in *Air*e^−/−^ mice. We found that selective JAK1 and JAK2 inhibition ameliorated autoimmune tissue damage and suppressed IFN-γ signaling, with JAK2 inhibition providing protection comparable to ruxolitinib. In contrast, selective JAK3 inhibition reduced lymphocyte accumulation but paradoxically increased the proportion of IFN-γ-producing T cells and failed to confer equivalent protection from autoimmunity. Together, these findings reveal a hierarchical organization of JAK signaling in AIRE deficiency, identify the IFN-γ-JAK2 axis as the dominant pathogenic pathway, and have important biological and translational implications for the rational design of targeted therapies in IFN-γ-mediated autoimmunity.

## Results

### Selective JAK inhibition decreases lymphocyte accumulation but differentially alters their parenchymal localization in the lung

CD4^+^ T cells are necessary and sufficient to drive autoimmunity in APECED [8, 9]. We therefore compared the effects of selective JAK1, JAK2, and JAK3 inhibition with those of the JAK1/2 inhibitor ruxolitinib on immune cell accumulation in *Air*e^−/−^ mice, using the lung as a sentinel tissue. The lung is uniformly affected by autoimmunity in *Air*e^−/−^ mice on the non-obese diabetic (NOD) background [17], in contrast to the variable involvement of other organs. Mice were treated with orally administered JAK inhibitors beginning at 3-5 weeks of age, a timeframe at which early autoimmune inflammation is already evident in the lung and other tissues, using a strategy previously described [12, 18].

Flow cytometric analysis showed that selective inhibition of JAK1, JAK2, or JAK3 resulted in a significant reduction in the total number of CD45^+^ leukocytes in the lungs of *Air*e^−/−^ mice as compared with untreated mice (Fig. 1A). Consistent with this finding, the numbers of TCRαβ^+^ T cells (Fig. 1B), CD4^+^ T cells (Fig. 1C), and CD8^+^ T cells (Fig. 1D) were decreased in the lungs of mice treated with each of the JAK inhibitors. In addition, TCRγδ^+^ T cell, innate lymphoid cell, and non-lymphoid immune cell populations were also reduced following selective inhibition of JAK1, JAK2, and JAK3 (Fig. 1−figure supplement 1). Thus, inhibition of JAK1, JAK2, or JAK3 broadly decreased leukocyte and lymphocyte accumulation in the lungs of *Air*e^−/−^ mice.

**Figure 1.**
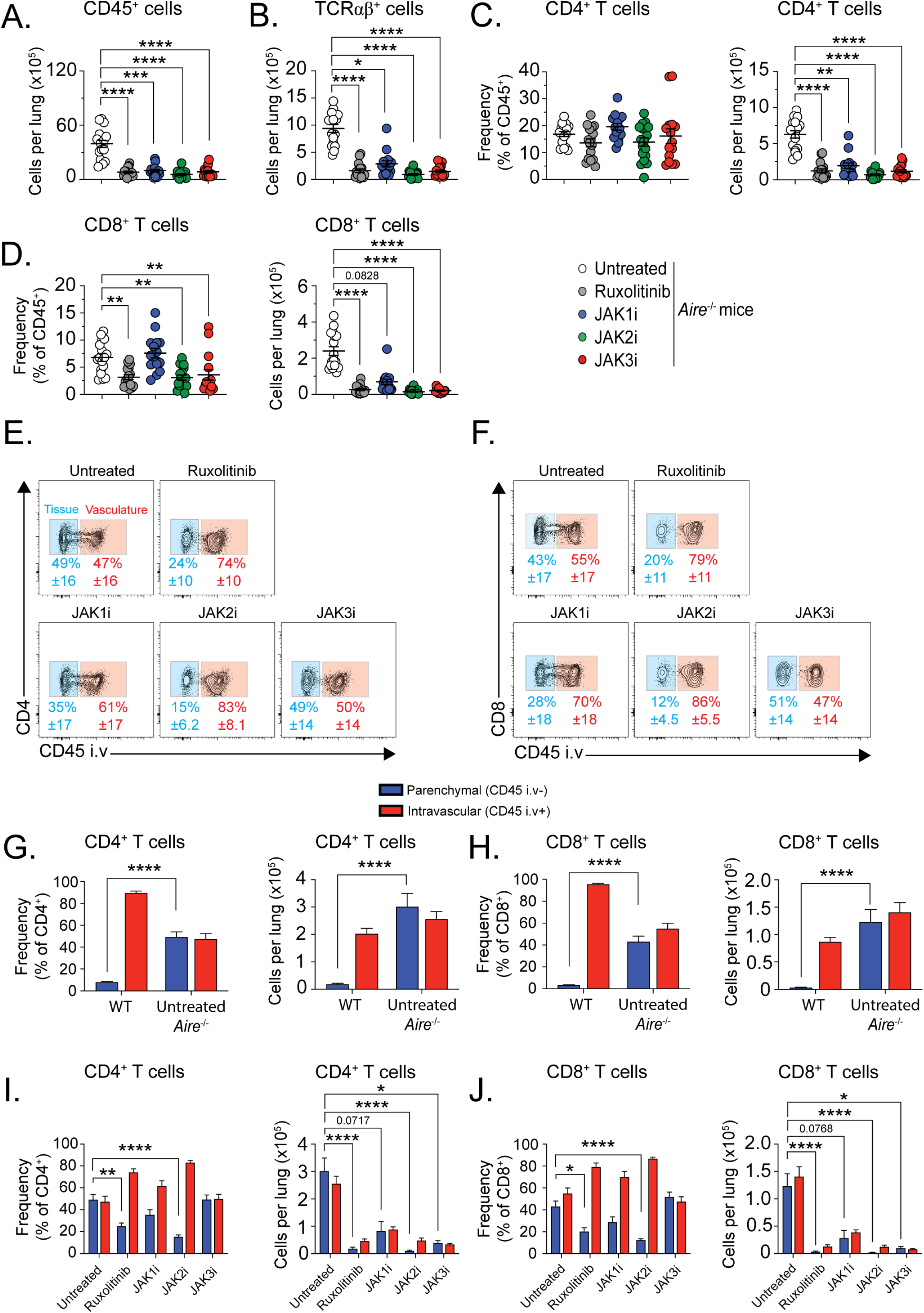
Selective JAK inhibition decreases lymphocyte accumulation but differentially alters their parenchymal localization in the lung. *Air*e^−/−^ mice were treated for four weeks with selective JAK inhibitors; untreated *Air*e^−/−^ and ruxolitinib-treated *Air*e^−/−^ mice served as controls. Lung lymphocytes were analyzed by flow cytometry. (**A**) Absolute numbers of CD45^+^ leukocytes and (**B**) TCRαβ^+^ T cells. (**C-D**) Frequency (of total CD45^+^) and absolute numbers of CD4^+^ and CD8^+^ T cells. (**E-F**) Representative flow cytometry plots showing parenchymal (CD45 i.v.^−^) or intravascular (CD45 i.v.^+^) CD4^+^ and CD8^+^ T cells. (**G-H**) Frequency and absolute numbers of parenchymal and intravascular CD4^+^ and CD8^+^ T cells in *Aire^+/+^*and untreated *Air*e^−/−^ *mice*. (**I-J**) Frequency and absolute numbers of parenchymal and intravascular CD4^+^ and CD8^+^ T cells in untreated and JAK inhibitor-treated *Air*e^−/−^ mice. For total CD45^+^, TCRαβ^+^, CD4^+^ T, and CD8^+^ T cells: *n* = 15-18 mice per group from four independent experiments. For parenchymal/intravascular ratio analyses: *n* = 11-15 mice per group from three independent experiments. Statistical comparisons between *Aire^+/+^* and untreated *Air*e^−/−^ mice were performed using the Mann-Whitney test. Comparisons among groups were performed using one-way ANOVA or Kruskal-Wallis test with Dunn’s multiple comparisons to untreated *Air*e^−/−^ mice. *P < 0.05, **P < 0.01, ***P < 0.001, ****P < 0.0001.

To determine whether selective JAK inhibition differentially affected the localization of T cells within the pulmonary parenchyma versus the intravascular compartment, we next assessed the relative distribution of intravascular and parenchymal CD4^+^ and CD8^+^ T cells using intravascular labeling with fluorescent anti-CD45 antibody immediately before euthanasia, as previously described (Fig. 1E-F) [19]. In *Air*e^−/−^ mice, there is a marked increase in the proportion of tissue-infiltrating, parenchymal (CD45 iv^−^) over intravascular (CD45 iv^+^) CD4^+^ and CD8^+^ T cells as compared with wild-type mice (Fig. 1G-H). Treatment with ruxolitinib or selective JAK2 inhibition significantly reduced the proportion of parenchymal CD4^+^ and CD8^+^ T cells and partially normalized the parenchymal-to-intravascular ratio relative to that observed in wild-type mice (Fig. 1I-J). In contrast, JAK1 or JAK3 inhibition did not significantly alter the proportion of parenchymal versus intravascular CD4^+^ and CD8^+^ T cells, despite reducing the overall number of T cells in the lung (Fig. 1I-J).

Taken together, these findings indicate that although inhibition of JAK1, JAK2, or JAK3 decreases total leukocyte and lymphocyte accumulation in the lungs of *Air*e^−/−^ mice, selective JAK inhibition exerts distinct effects on T cell tissue localization, with ruxolitinib JAK2 inhibition reversing the enhanced parenchymal accumulation feature of AIRE deficiency.

### Selective JAK3 inhibition uncouples lymphocyte burden from IFN-γ-dominant signaling

Given the pathogenic role of IFN-γ in driving autoimmune tissue injury in APECED, we next examined whether the distinct effects of selective JAK inhibition on lymphocyte accumulation and tissue localization were associated with differential regulation of IFN-γ production and downstream signaling in the lung. We therefore assessed IFN-γ production by CD4^+^ and CD8^+^ T cells, as well as IFN-γ-responsive transcriptional and protein readouts, in *Air*e^−/−^ mice treated with selective JAK1, JAK2, or JAK3 inhibitors or with ruxolitinib.

Flow cytometric analysis revealed a striking divergence between lymphocyte burden and cytokine bias following selective JAK inhibition (Fig. 2A-B). Selective JAK3 inhibition resulted in a paradoxical increase in the proportion of IFN-γ-producing CD4^+^ and CD8^+^ T cells in the lung compared with untreated mice and with mice treated with ruxolitinib, JAK1 inhibition, or JAK2 inhibition (Fig. 2C-D). Consequently, the absolute number of IFN-γ-producing CD4^+^ T cells was not significantly reduced in JAK3 inhibitor-treated mice relative to other treatment groups, despite a reduction in total T cell numbers. In contrast, the most pronounced reductions in the numbers of IFN-γ-producing CD4^+^ and CD8^+^ T cells were observed following treatment with ruxolitinib or selective JAK2 inhibition, whereas JAK1 inhibition exerted more modest, intermediate effects (Fig. 2C-D).

**Figure 2.**
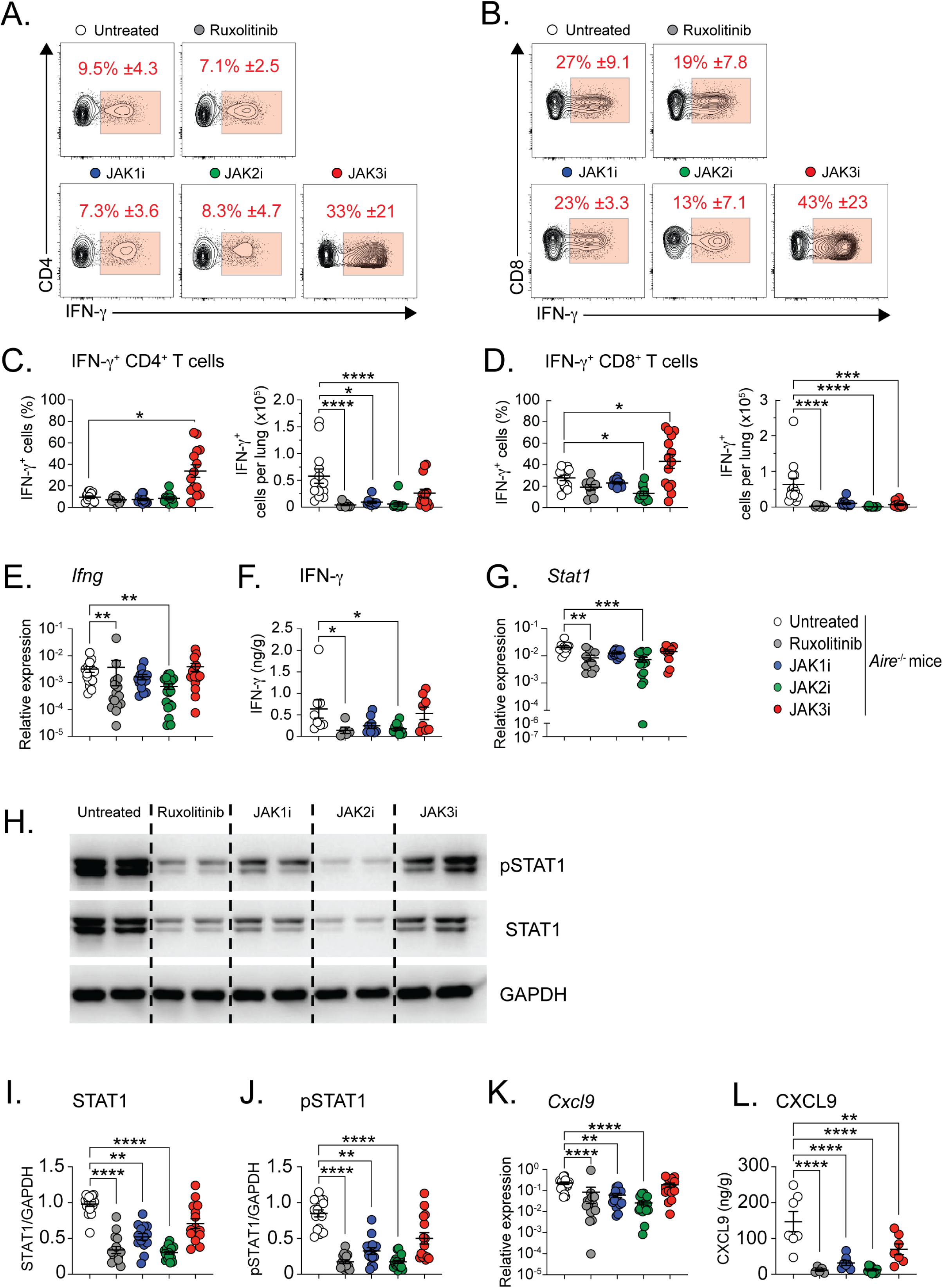
Selective JAK3 inhibition uncouples lymphocyte burden from IFN-γ-dominant signaling. Untreated *Air*e^−/−^ mice and *Air*e^−/−^ mice treated for four weeks with selective JAK1i, JAK2i, or JAK3i were analyzed. Lungs were processed for intracellular cytokine staining, qPCR, ELISA, and immunoblot analyses. (**A-B**) Representative flow cytometry plots showing IFN-γ production by CD4^+^ and CD8^+^ T cells. (**C-D**) Frequency (of total CD4^+^ and CD8^+^ T cells) and absolute numbers of IFN-γ^+^CD4^+^ and IFN-γ^+^CD8^+^ T cells in the lung. (**E-F**) Relative *Ifng* mRNA expression and IFN-γ protein concentrations in lung homogenates. (**G**) Relative *Stat1* mRNA expression. (**H**) Representative immunoblots of phospho-STAT1 (pSTAT1), total STAT1, and GAPDH. (**I-J**) Quantification of total STAT1 and phospho-STAT1 normalized to GAPDH. For IFN- γ^+^CD4^+^ and IFN-γ^+^CD8^+^ T cells: *n* = 9-14 mice per group from four independent experiments. For *Ifng* and *Cxcl9* mRNA: *n* = 15-22 mice per group from four independent experiments. For *Stat1*_mRNA: *n* = 10-17 mice per group from three independent experiments. For CXCL9 protein: *n* = 5-10 mice per group from two independent experiments. For STAT1 and pSTAT1 immunoblots: *n* = 15-22 mice per group from four independent experiments). Statistical analyses were performed using one-way ANOVA or Kruskal-Wallis test with Dunn’s multiple comparisons to the untreated *Air*e^−/−^ mice. *P < 0.05, **P < 0.01, ***P < 0.001, ****P < 0.0001.

Consistent with these findings, levels of *Ifng* mRNA (Fig. 2E) and IFN-γ protein (Fig. 2F) in lung tissue were significantly decreased in mice treated with ruxolitinib or selective JAK2 inhibition but not in mice treated with JAK1 or JAK3 inhibition. Downstream of IFN-γ receptor engagement, ruxolitinib and selective JAK2 inhibition caused the greatest suppression of *Stat1* mRNA expression (Fig. 2G), and total STAT1 protein levels as assessed by immunoblot analysis (Fig. 2H-I), whereas selective JAK1 inhibition had lesser effects and selective JAK3 inhibition exerted no significant effect. Phosphorylation of STAT1 was similarly reduced by ruxolitinib and selective JAK2 inhibition, with a smaller but significant reduction observed with selective JAK1 inhibition and no significant change following selective JAK3 inhibition (Fig. 2J). Expression of CXCL9, a canonical IFN-γ-inducible chemokine, mirrored this hierarchy. Both *Cxcl9* mRNA (Fig. 2K) and CXCL9 protein levels (Fig. 2L) were more profoundly reduced by ruxolitinib and selective JAK2 inhibition, followed by a lesser decrease with selective JAK1 inhibition, whereas selective JAK3 inhibition resulted in mild suppression at the protein level.

Taken together, these findings demonstrate that selective JAK inhibition differentially regulates IFN-γ-dominant immune programs in AIRE deficiency. Notably, selective JAK3 inhibition uncouples reductions in lymphocyte accumulation from suppression of IFN-γ production and signaling, whereas ruxolitinib and selective JAK2 inhibition most effectively suppress IFN-γ-driven pathogenic pathways in the lung.

### JAK2 inhibition most effectively ameliorates autoimmune tissue injury

Given the differential effects of selective JAK inhibition on lymphocyte accumulation, tissue localization, and IFN-γ-dominant signaling in the lung, we next examined whether these immunological differences translated into corresponding effects on autoimmune tissue injury. To quantify disease severity in an unbiased manner, we employed an image analysis platform to determine the percentage of inflamed or damaged tissue in histologic sections (Fig. 3−figure supplement 1), expressing treatment effects as residual inflammation relative to untreated *Air*e^−/−^ mice.

**Figure 3.**
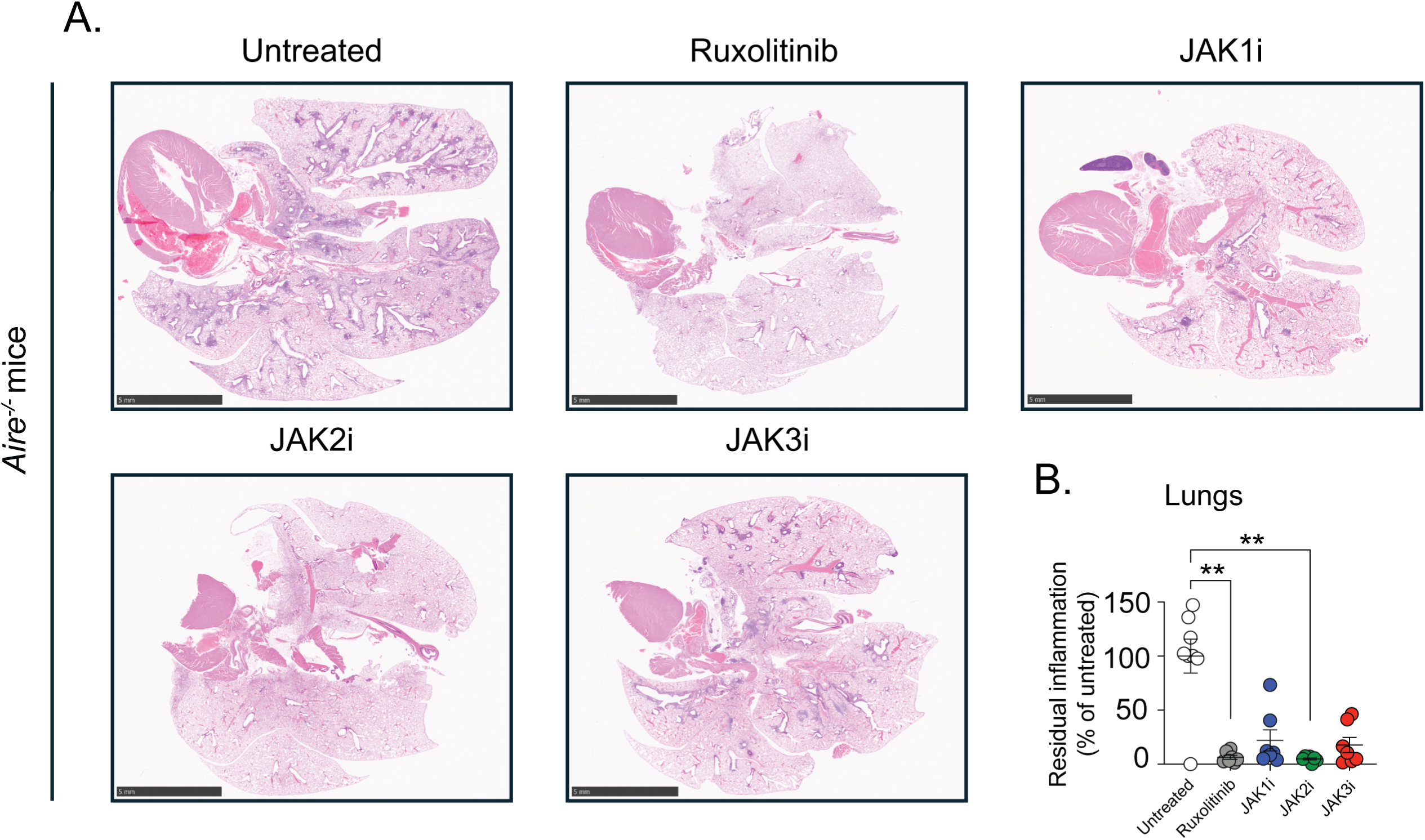
JAK2 inhibition most effectively ameliorates autoimmune tissue injury in the lungs. Untreated *Air*e^−/−^ mice and *Air*e^−/−^ mice treated for four weeks with selective JAK1i, JAK2i, or JAK3i were analyzed. Lungs were harvested and processed for histopathological assessment. (**A**) Representative hematoxylin and eosin (H&E)-stained sections of lung (0.5x; scale bar = 5 mm) from untreated *Air*e^−/−^ mice and JAK inhibitor-treated *Air*e^−/−^ mice. (**B**) Quantification of residual tissue inflammation in lungs expressed as percentage of inflammation relative to untreated *Air*e^−/−^ mice. All samples were normalized to untreated *Air*e^−/−^ controls within each independent experiment. *n* = 7-8 mice per group from 3 independent experiments. Statistical comparisons were performed using Kruskal-Wallis test with Dunn’s multiple comparisons to the untreated *Air*e^−/−^ mice. *P < 0.05, **P < 0.01, ***P < 0.001, ****P < 0.0001.

Consistent with their pronounced effects on T cell accumulation and localization and IFN-γ signaling, treatment with ruxolitinib or selective JAK2 inhibition resulted in the most robust protection from autoimmune lung injury, with greater than 90% average reduction in histologic inflammation compared with untreated mice (Fig. 3). In contrast, selective JAK1 and JAK3 inhibition conferred partial protection with less improvement in lung pathology (Fig. 3).

We next extended these analyses to two additional organs commonly affected by autoimmunity in *Air*e^−/−^ mice, the salivary glands and eyes, which exhibit autoimmune involvement in most, but not all, animals. In both tissues, ruxolitinib and selective JAK2 inhibition again produced the most substantial improvement in histological abnormalities with greater than 90% average reduction in tissue injury (Figs. 4 and 5). Selective JAK1 inhibition resulted in more modest improvement, with approximately 40-60% average reduction in histologic damage, whereas selective JAK3 inhibition was least effective, conferring less than 30-40% average improvement in both organs, an effect that did not reach statistical significance.

**Figure 4.**
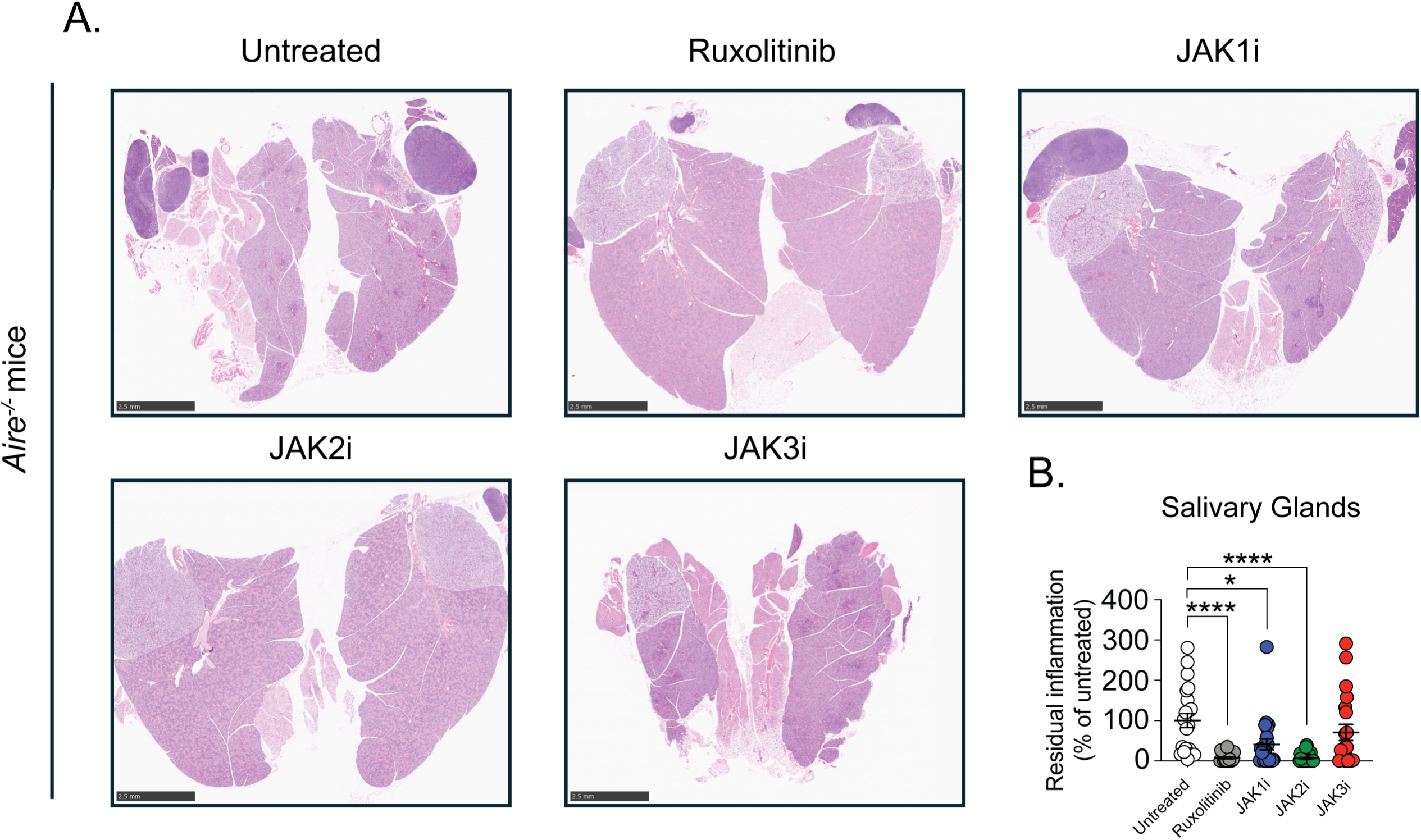
JAK2 inhibition most effectively ameliorates autoimmune tissue injury in the salivary glands. Untreated *Air*e^−/−^ mice and *Air*e^−/−^ mice treated for four weeks with selective JAK1i, JAK2i, or JAK3i were analyzed. Salivary glands were harvested and processed for histopathological assessment. (**A**) Representative hematoxylin and eosin (H&E)-stained sections of salivary glands (0.75x; scale bar = 2.5 mm) from untreated *Air*e^−/−^ mice and JAK inhibitor-treated *Air*e^−/−^ mice. (**B**) Quantification of residual tissue inflammation in salivary glands expressed as percentage of inflammation relative to untreated *Air*e^−/−^ mice. All samples were normalized to untreated *Air*e^−/−^ controls within each independent experiment. *n* = 17-25 mice per group from six independent experiments. Statistical comparisons were performed using Kruskal-Wallis test with Dunn’s multiple comparisons to the untreated *Air*e^−/−^ mice. *P < 0.05, **P < 0.01, ***P < 0.001, ****P < 0.0001.

**Figure 5.**
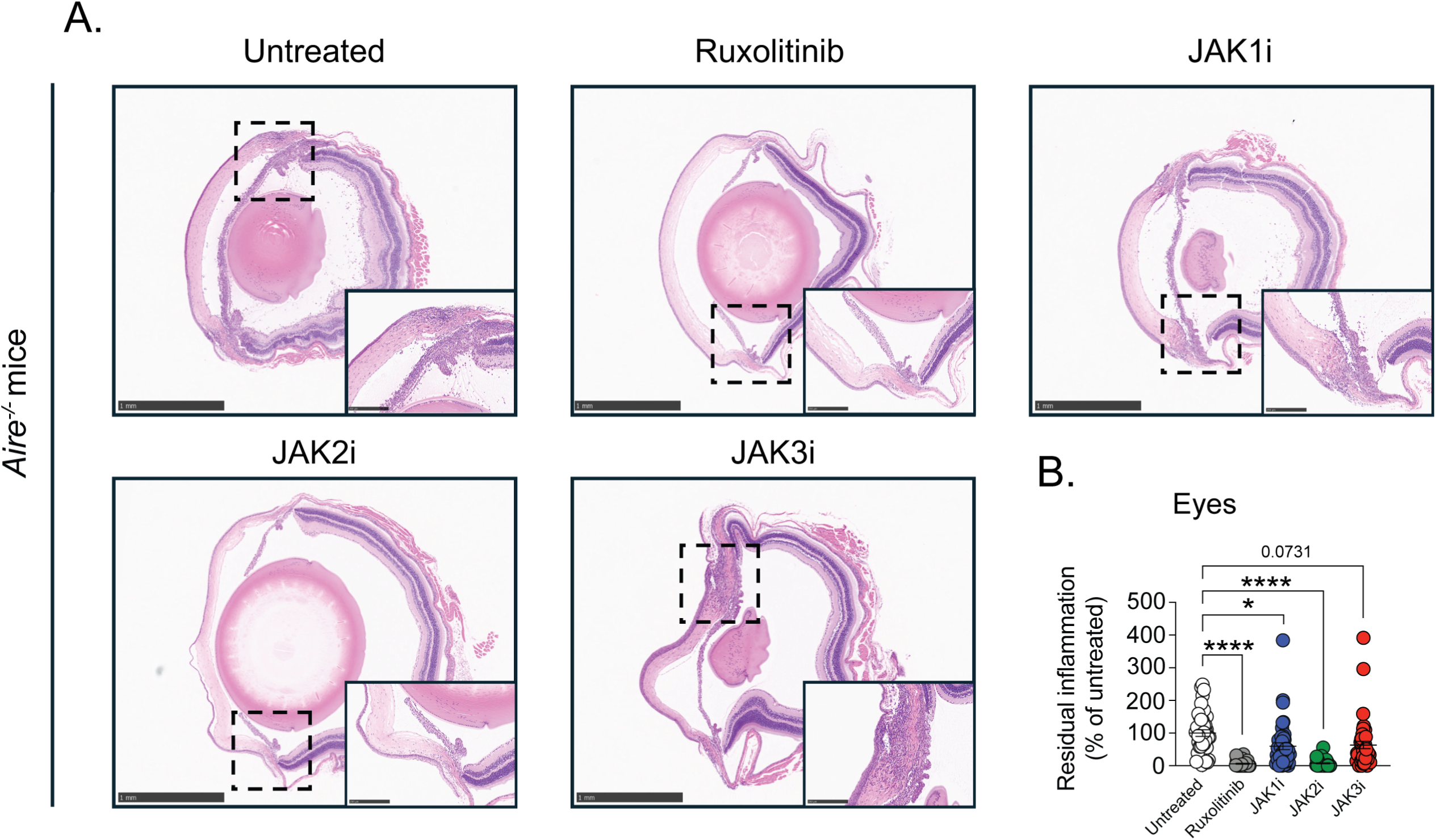
JAK2 inhibition most effectively ameliorates autoimmune tissue injury in the eyes. Untreated *Air*e^−/−^ mice and *Air*e^−/−^ mice treated for four weeks with selective JAK1i, JAK2i, or JAK3i were analyzed. Eyes were harvested and processed for histopathological assessment. (**A**) Representative hematoxylin and eosin (H&E)-stained sections of eyes (2.5x; scale bar = 1 mm) from untreated *Air*e^−/−^ mice and JAK inhibitor-treated *Air*e^−/−^ mice. The insets provide a greater magnification of the ciliary body area of the inflamed eye (10x; scale bar = 250 µm). (**B**) Quantification of residual tissue inflammation in eyes expressed as percentage of inflammation relative to untreated *Air*e^−/−^ mice. All samples were normalized to untreated *Air*e^−/−^ controls within each independent experiment. *n* = 17-25 mice per group from six independent experiments. Statistical comparisons were performed using Kruskal-Wallis test with Dunn’s multiple comparisons to the untreated *Air*e^−/−^ mice. *P < 0.05, **P < 0.01, ***P < 0.001, ****P < 0.0001.

Taken together, these data demonstrate that selective JAK inhibition differentially impacts autoimmune tissue injury in *Air*e^−/−^ mice and identify JAK2 inhibition as the most effective strategy for ameliorating multiorgan autoimmunity, with efficacy comparable to that of the nonselective JAK1/2 inhibitor, ruxolitinib.

## Discussion

In this study, we define a hierarchical organization of JAK signaling in AIRE deficiency and identify IFN-γ-JAK2 signaling as the dominant pathogenic axis during autoimmune tissue injury. Although selective inhibition of JAK1, JAK2, or JAK3 reduced overall lymphocyte accumulation in affected tissues, only ruxolitinib and selective JAK2 inhibition effectively suppressed IFN-γ production, downstream STAT1 signaling, and multiorgan autoimmune pathology. In contrast, JAK3 inhibition uncoupled reductions in lymphocyte burden from control of pathogenic cytokine programs, paradoxically enriching IFN-γ-producing T cells and failing to significantly ameliorate tissue injury. These findings demonstrate that lymphocyte accumulation and cytokine bias are separable features of autoimmune inflammation in AIRE deficiency and establish IFN-γ signaling, rather than generalized lymphocyte expansion, as a principal determinant of disease severity.

A particularly unexpected observation was the paradoxical enrichment of IFN-γ-producing T cells following selective JAK3 inhibition. JAK3 mediates signaling downstream of γc cytokines, including IL-2, IL-4, IL-7, IL-9, IL-15, and IL-21, which play central roles in T cell survival, differentiation, and homeostasis [20]. The exacerbation of IFN-γ bias observed upon JAK3 inhibition raises the possibility that γc cytokine signaling may restrain the IFN-γ-dominant pathogenic program in the setting of AIRE deficiency, revealing a potential layer of cross-regulatory control that becomes unmasked when γc signaling is selectively disrupted. Although the precise mechanisms underlying this effect remain to be defined, these findings highlight an important principle: suppression of lymphocyte survival or expansion does not necessarily equate to suppression of pathogenic cytokine programs and may, in certain settings, skew immune responses toward increased inflammatory potential.

Our results also underscore marked tissue-specific and context-specific differences in JAK pathway dominance across IFN-γ-mediated diseases. In alopecia areata, an IFN-γ-associated inflammatory condition of the skin, selective JAK3 inhibition –now FDA-approved for clinical use– has been reported to exert the strongest therapeutic effects, followed by JAK1 inhibition, with relatively limited benefit from JAK2 inhibition in preclinical models [14]. In striking contrast, we find that in AIRE deficiency, JAK2 inhibition phenocopies the protective effects of broad JAK1/2 blockade, whereas JAK3 inhibition is ineffective. These opposing hierarchies suggest that IFN-γ-driven inflammation is not governed by a uniform JAK logic across tissues and settings but instead reflects context-dependent integration of cytokine networks, cellular composition, and tissue microenvironment.

The dominance of JAK2 in AIRE deficiency is mechanistically consistent with the central role of IFN-γ signaling in this disease. Our findings that selective JAK2 inhibition suppresses IFN-γ production, STAT1 activation, IFN-γ-inducible chemokine expression, and tissue injury place JAK2 at the apex of pathogenic signaling in AIRE deficiency. By contrast, selective JAK1 inhibition provided intermediate effects, whereas selective JAK3 inhibition failed to suppress the IFN-γ axis despite reducing lymphocyte numbers. Although IFN-γ signals through a receptor complex that engages both JAK1 and JAK2, prior studies have demonstrated differential dependence on these kinases for IFN-γ-induced transcriptional programs and STAT1 activation. Notably, IFN-γ-dependent gene expression and STAT1 phosphorylation can be dominantly JAK2-dependent, with kinase-inactive or deficient JAK1 –but not JAK2– capable of sustaining IFN-γ-responsive gene expression despite impaired antiviral signaling [21, 22], providing a potential mechanistic framework for the more limited efficacy of selective JAK1 inhibition observed in our study.

These observations have important translational implications. Broad JAK inhibition, including with ruxolitinib, has demonstrated clinical efficacy in IFN-γ-mediated diseases but can be associated with well-recognized toxicities, including increased susceptibility to viral infections, disruption of lymphocyte homeostasis, thrombotic events, and malignancies [15, 16]. Our data suggest that selective targeting of JAK2 may preserve therapeutic efficacy in IFN-γ-driven autoimmunity while sparing JAK1- and γc-dependent immune programs that are critical for host defense and immune homeostasis. Although these findings do not directly address long-term safety, they provide a mechanistic rationale for considering JAK2-selective strategies to narrow the therapeutic window of JAK inhibition.

Our study has several limitations. Our experiments were performed in *Air*e^−/−^ mice, and although this model recapitulates key immunological and pathologic features of human APECED [23], species-specific differences in immune regulation may influence the relative contribution of individual JAK pathways. In addition, the mechanisms underlying the paradoxical enrichment of IFN-γ-producing T cells following selective JAK3 inhibition were not directly examined and warrant further investigation. Finally, differences in pharmacokinetic and pharmacodynamic properties among the JAK inhibitors used could contribute to their differential effects observed *in vivo*. We did not directly measure drug concentrations or target engagement in tissues, and comprehensive *in vivo* pharmacokinetic analysis, such as drug level quantification by LC-MS/MS, are technically challenging and not routinely feasible for these compounds. As such, although our findings define clear biological hierarchies of JAK pathway dependence, future studies integrating pharmacologic and mechanistic approaches will be important to further refine these conclusions.

In summary, our findings reveal a hierarchical organization of JAK signaling in AIRE deficiency, identify IFN-γ-JAK2 signaling as the dominant driver of autoimmune tissue injury, and demonstrate that suppression of lymphocyte accumulation alone is insufficient to control disease. By distinguishing pathogenic cytokine signaling from generalized immune suppression, this work provides a conceptual framework for rational, pathway-directed immunomodulation in IFN-γ-mediated autoimmunity.

## Methods

### Sex as biological variable

We used both male and female animals and found similar results. Accordingly, all data represent combined analyses of male and female mice from three to four independent experiments.

### Animals

Non-obese Diabetic (NOD) *Aire*^+/−^ mice were purchased from The Jackson Laboratory (strain no. 006360) and bred as heterozygous pairs to attain *Aire*^−/−^ and *Aire*^+/+^ littermate mice for experiments. Breeders and their progeny were housed in NIH animal facilities. Both female and male mice were used in experiments and treatments were initiated at 3-5 weeks of age. All mouse experiments were performed according to guidelines set forth by the Guide for the Care and Use of Laboratory Animals under a protocol approved by the NIAID Animal Care and Use Committee (protocol: LCIM14E).

### Treatment with selective JAK inhibitors

Ruxolitinib phosphate (HY-50858), itacitinib (HY-16997), CEP-33779 (HY-15343), and ritlecitinib (HY-100754) were used as selective inhibitors for JAK1/2, JAK1, JAK2, and JAK3, respectively, as previously reported [14]. Selective inhibitors for JAK1, JAK2, and JAK3 are referred to as JAK1i, JAK2i, and JAK3i in the figures of the manuscript. All inhibitors were purchased from MedChemExpress.

Nutra-Gel Diet^TM^ bacon flavored feed (catalog no. F5769-KIT; Bio-Serv, USA) was used for all experiments and prepared according to the manufacturer’s instructions. Before treatment initiation, mice were acclimated to the diet without drug for 2-3 days. Mice were then treated with their respective JAK inhibitors at a dosage of 0.4 g of drug per 1 kilogram of food. Selective JAK1, JAK2, and JAK3 inhibitors were dissolved in 100% DMSO to a concentration of 11.2 mg/mL and subsequently mixed into the feed. Control feed for untreated *Aire^+/+^* and *Aire*^−/−^ mice was mixed with an equivalent concentration of DMSO (4.84% DMSO per tray of food). In contrast, ruxolitinib –used as a positive control– was water soluble administered at a dosage of 1.0 g of drug per 1 kilogram of food. Each treatment group was distinguished by food coloring, and fresh food provided daily for four weeks.

### Histopathology

Mouse eyes, salivary glands, and lungs were harvested and fixed in 10% buffered formalin for 48 hours, then transferred to 70% ethanol and processed for hematoxylin and eosin (H&E) staining (Histoserv, Inc., USA). Slides were digitally scanned, and areas of inflammation were quantified. QuPath image analysis software was used for lung sections, where pixel thresholds were generated to identify stromal regions and areas of inflammation. A representative example of the annotation is shown in Figure 3−figure supplement 1. Motic histology viewing software was used to manually quantify inflammatory areas in eye and salivary gland sections, as these organs were not compatible with automated QuPath analysis. Areas of inflammation on the histologic sections were scored in a blinded manner as above and the treatment effects were quantified as residual inflammation relative to untreated *Air*e^−/−^ mice.

### RNA extraction and qPCR

Mouse lung tissue was placed in TRIzol reagent (catalog no. 15596026; Invitrogen, USA), homogenized with 5 mm stainless steel beads (catalog no. 69989; Qiagen, USA) using a TissueLyser II (Qiagen, USA), centrifuged at 16,000 x g for 10 minutes at 4°C, and stored at -80°C until RNA extraction. RNA was extracted using the RNeasy Mini Kit (catalog no.74106; Qiagen, USA), and RNA concentration was determined using a NanoDrop One spectrophotometer (Thermo Fisher Scientific, USA). cDNA was synthesized with 1 µg of RNA using the qScript^TM^ cDNA synthesis kit (catalog no. 95047-500; Quantabio, USA). qPCR was performed with 100 ng of cDNA using TaqMan^TM^ reagents on a QuantStudio^TM^ 3 system. The TaqMan^TM^ probes used were *Gapdh* (Mm99999915_g1), *Ifng* (Mm01168134_m1), *Cxcl9* (Mm00434946_m1), and *Stat1* (Mm01257286_m1). All samples were normalized to *Gapdh,* and results were presented on a logarithmic scale.

### ELISA

Mouse lung tissue was placed in ELISA buffer, grinded with 5 mm stainless steel beads in a TissueLyser II, centrifuged at 16,000 x g for 10 minutes at 4°C, and stored at -80°C until analysis. ELISA buffer consisted of PBS supplemented with 0.05% Tween-20 and cOmplete™, Mini, EDTA-free Protease Inhibitor Cocktail (catalog no. 11836170001; Roche, USA). A DuoSet® ELISA kit (catalog no. DY492-05; R&D Systems, USA) was used to measure CXCL9 and a high-sensitivity ELISA kit was used for IFN-γ (catalog no. 88-8314-88; Invitrogen, USA). All assays were performed according to manufacturer’s instructions.

### Western Blot

A RIPA buffer cocktail was prepared using RIPA Lysis and Extraction Buffer (catalog no. 89901; ThermoFisher Scientific, USA), Halt™ Protease Inhibitor Cocktail (catalog no. 87786; ThermoFisher Scientific, USA), and Halt™ Phosphatase Inhibitor Single-Use Cocktail (catalog no. 78420; ThermoFisher Scientific, USA). Mouse lung tissue was homogenized in RIPA buffer, homogenized with 5 mm stainless steel beads in a TissueLyser II, centrifuged at 16,000 x g for 10 minutes at 4°C, and stored at -80°C. Protein concentration was determined using the Bradford assay (Bio-Rad, USA). Proteins were resolved on 10% SDS-PAGE gels and were transferred to 0.2 µm PVDF membranes using a Trans-Blot Turbo^TM^ system (Bio-Rad, USA). Membranes were blocked with 5% nonfat milk in TBS containing 0.1% Tween-20 for 1 hour at room temperature, washed with 0.1% Tween-20 in TBS, and incubated with primary antibodies overnight at 4°C. Membrane was washed again three times and incubated with secondary antibodies for 1 hour at room temperature. Blots were developed using Clarity^TM^ Western ECL (catalog no. 1705061; Bio-Rad, USA) or Radiance Plus Femtogram HRP substrates (catalog no. AC2103; Azure Biosystems, USA) and imaged using a ChemiDoc^TM^ system (Bio-Rad, USA). Densitometric analysis was performed using FIJI software. Primary antibodies against GAPDH (catalog no. 5174S), STAT1 (catalog no. 14995S), and pSTAT1 (catalog no. 9167S) (Cell Signaling Technologies, USA) were used at 1:1,000 dilution, with HRP-conjugated anti-rabbit IgG conjugated with HRP (catalog no.7074S; Cell Signaling Technology) as the secondary antibody.

### Intravascular staining and lung digestion

Mice were injected intravenously with 3 μg AlexaFluor 488-conjugated anti-mouse CD45 antibody (catalog no.103122; BioLegend, USA). Three minutes later, lungs were harvested and placed in gentleMACS^TM^ C tubes (catalog no.130-093-237; Miltenyi Biotec, Germany) containing 1 mg/ml collagenase IV (catalog no. LS004189; Worthington Biochemical Corporation, USA) and 20 μg/ml DNase I (catalog no. DN25-1G; Millipore-Sigma, USA) in RPMI-1640. Tissue was mechanically dissociated in gentleMACS^TM^ Octo Dissociator (Miltenyi Biotec, Germany) with program “mlung 01_02”, incubated for 25 minutes at 37°C, and further dissociated using program “mlung 02_01”. Cell suspensions were resuspended in 40% Percoll (catalog no. 17-0891-01; Cytiva, USA), underlaid with 66% Percoll, and centrifuged for 20 minutes at 805 x g at room temperature. The interphase was collected, filtered through a 40 μm strainer, and resuspended in complete RPMI containing 10% FBS and penicillin/streptomycin.

### Flow cytometry

Single cell suspensions were stained with LIVE/DEAD Fixable Blue Dead Cell Stain kit (catalog no. L23105; Invitrogen, USA) and Fc Block (catalog no. 553141; BD, USA) for 10 minutes at room temperature, washed, and incubated with the following antibodies (all from BioLegend, USA) for 30 minutes at 4°C: PE-Cy7-conjugated anti-mouse TCRγ (catalog no. 25-5711-82; Invitrogen, USA), BV421-conjugated anti-mouse TCRβ (catalog no. 109229), BV510-conjugated anti-mouse CD4 (catalog no. 100559), AlexaFluor 700-conjugated anti-mouse CD8 (catalog no. 100730), BV650-conjugated anti-mouse CD44 (catalog no. 103049), BV785-conjugated anti-mouse CD45 (catalog no. 103149), and APC-Fire750-conjugated anti-mouse CD90.2 (catalog no. 140326). Cells were fixed in 2% formaldehyde and Spherotec Counting Beads (catalog no. ACFP-70-10; Spherotec Inc., USA) were added to determine the absolute cell numbers. Samples were acquired on an LSR Fortessa, and the data analyzed using FlowJo v10.10.0 (BD, USA).

### T cell stimulation and intracellular cytokine staining

For *ex vivo* stimulation, lung cell suspensions were incubated for 4 hours at 37°C with 50 ng/ml PMA (catalog no. 16561-29-8; Millipore Sigma, USA), 2.5 μg/ml ionomycin (catalog no. I9657; Millipore Sigma), 1 μg/ml brefeldin A (catalog no. 555029; BD, USA) and 2 μM monensin (catalog no. 420701; BioLegend, USA) in RPMI-1640 containing penicillin/streptomycin and 10% heat-inactivated fetal bovine serum. Four hours later, cells were surface-stained, fixed, permeabilizated with fixation/permeabilization solution (catalog no. 51-2090KZ; BD, USA) for 20 minutes, washed with perm/wash buffer (Cat.51-2091KZ, BD, USA), and stained intracellularly with PE-conjugated anti-mouse IFN-γ (catalog no. 505808; BioLegend, USA). Samples were acquired on an LSR Fortessa and analyzed using FlowJo.

### Statistics

Statistical analyses were performed sing GraphPad Prism 10. Data normality was assessed using the Shapiro-Wilk test. If all groups passed normality testing, a one-way ANOVA parametric test with Dunnett’s multiple comparisons to the untreated control group was used. If any group failed the normality test, a Kruskal-Wallis test with Dunn’s multiple comparisons to the untreated control group was applied. Statistical comparisons between *Aire^+/+^* and untreated *Air*e^−/−^ mice in figure 1G-H were performed using the Mann-Whitney test.

## Data availability

All data analyzed during this study are included in the manuscript and the accompanying supplementary materials. This includes all primary quantitative data underlying figures and tables, complete descriptions of experimental conditions, and all observations necessary to reproduce the reported findings. No custom code, sequencing data, or large-scale omics datasets were generated in this study. Data sharing complies with applicable institutional, legal, and ethical standards.

## Authors contributions

EH and LDSD performed experiments and analyzed the data. EH, LDSD, and MSL designed the experiments and wrote the paper. MSL conceived and supervised the execution of the project.

## Acknowledgments

The authors thank Andew L. Wishart for his technical assistance in harvesting some mouse samples. This research was supported by the Intramural Research Program of the National Institutes of Health (NIH). The contributions of the NIH authors were made as part of their official duties as NIH federal employees, are in compliance with agency policy requirements, and are considered Works of the United States Government. However, the findings and conclusions presented in this paper are those of the authors and do not necessarily reflect the views of the NIH or the U.S. Department of Health and Human Services.

## Supplemental Figure Legends

**Figure 1−figure supplement 1. Selective JAK inhibition decreases TCRγδ^+^ T cell, innate lymphoid cell, and non-lymphoid cell accumulation in the lung.** *Air*e^−/−^ mice were treated for four weeks with selective JAK inhibitors; untreated *Air*e^−/−^ and ruxolitinib-treated *Air*e^−/−^ mice served as controls. Lung lymphocytes were analyzed by flow cytometry. (**A**) Absolute numbers of TCRγδ^+^ T cells, (**B**) innate lymphoid cells (CD90.2^+^TCRβ^−^, TCRγ^−^), and (**C**) non-lymphoid immune cells (CD45^+^CD90.2^−^). *n* = 15-18 mice per group from four independent experiments. Statistical comparisons were performed using Kruskal-Wallis test with Dunn’s multiple comparisons to the untreated *Air*e^−/−^ mice. *P < 0.05, **P < 0.01, ***P < 0.001, ****P < 0.0001.

**Figure 3−figure supplement 1. Histopathological analysis of lungs with QuPath.** Hematoxylin and eosin (H&E)-stained sections of lungs from all groups were scanned, and a pixel threshold for the outline of lung tissue and areas of inflammation was created in QuPath. Panel A shows a representative lung section stained with H&E before pixel thresholds were applied. Panel B shows an inset of panel A before analysis. Panel C shows the inset with annotations created with the pixel thresholds for lung tissue (green outline) and areas of inflammation (black outline).

